# Receptor for Hyaluronan-Mediated Motility (RHAMM) defines an invasive niche associated with tumor progression and predicts poor outcomes in breast cancer patients

**DOI:** 10.1101/2022.06.13.495375

**Authors:** Sarah E. Tarullo, Yuyu He, Claire Daughters, Todd P. Knutson, Christine Henzler, Matthew Price, Ryan Shanley, Patrice Witschen, Cornelia Tolg, Rachael E. Kaspar, Caroline Hallstrom, Lyubov Gittsovich, Megan L. Sulciner, Xihong Zhang, Colleen Forester, Oleg Shats, Michelle M. Desler, Kenneth Cowan, Douglas Yee, Kathryn L. Schwertfeger, Eva Turley, James B. McCarthy, Andrew C. Nelson

## Abstract

Breast cancer invasion and metastasis result from a complex interplay between tumor cells and the tumor microenvironment (TME). Key oncogenic changes in the TME include aberrant metabolism and subsequent signaling of hyaluronan (HA). Hyaluronan Mediated Motility Receptor (RHAMM, *HMMR*) is a HA receptor that enables tumor cells to sense and respond to the TME during breast cancer progression. Focused gene expression analysis of an internal breast cancer patient cohort demonstrates increased *RHAMM* expression correlates with aggressive clinicopathological features. We also develop a 27-gene RHAMM-dependent signature (RDS) by intersecting differentially expressed genes in lymph node positive cases with the transcriptome of a RHAMM-dependent model of cell transformation, which we validate in an independent cohort. We demonstrate RDS predicts for poor survival and associates with invasive pathways. Further analyses using CRISPR/Cas9 generated *RHAMM* -/- breast cancer cells provide direct evidence that RHAMM promotes invasion *in vitro* and *in vivo*. Additional immunohistochemistry studies highlight heterogeneous RHAMM expression, and spatial transcriptomics confirms the RDS emanates from RHAMM-high invasive niches. We conclude RHAMM upregulation leads to the formation of ‘invasive niches’, which are enriched in RDS-related pathways that drive invasion and could be targeted to limit invasive progression and improve patient outcomes.

## INTRODUCTION

Despite improvements in screening and treatment modalities, breast cancer remains the second leading cause of cancer-related deaths in women in the United States (1). Breast cancer mortality is impacted both by advanced stage disease at presentation, such as local metastasis to the lymph node, and by breast cancer recurrence (2). The capability of cancer cells to modify and infiltrate adjacent tissue is a hallmark common to all stages of breast cancer progression from initial invasion to distant metastasis (3). Mechanisms that drive breast cancer invasion and metastasis involve an intricate interplay between tumor cells and various aspects of the tumor microenvironment (TME).

Tumor-induced evolution of the TME is complex but characterized in part by active remodeling of the extracellular matrix (ECM) (4). A major component of the ECM is the glycosaminoglycan hyaluronan (HA), which is synthesized and modified both by tumor cells and cells within the TME (5, 6). HA synthesis and metabolism into low molecular weight fragments and oligomers are tightly regulated during homeostasis where HA primarily exists as a high molecular weight polymer, which has been shown to restrict neoplastic transformation (7). Subsequent detection of both low- and high-molecular weight HA by HA receptors, which include CD44, LYVE1, TLR2,4, and RHAMM, are a critical part of an early danger-sensing mechanism that detects tissue injury and that initiates cell responses needed for rapid repair (8-18). This injury detection mechanism is normally dampened once normal tissues are repaired but is hijacked by tumors to chronically maintain elevated HA production and metabolism, which facilitates malignant progression (6, 11). Tumor-induced dysregulation of HA metabolism includes not only increased synthesis of high molecular weight HA by one or more hyaluronan synthases (HAS1-3), but also increased enzymatic (HYAL1-2, CEMIP) or chemically (e.g. reactive oxygen species)-induced degradation to low molecular weight HA and oligomers (11-18). These low molecular weight HA fragments activate signaling pathways via HA receptors that support tumor proliferation, survival, invasion, and metastasis in many cancer types (6, 19, 20). Although studies have historically focused on HA as an ECM component, recent evidence has implicated intracellular HA as important for regulating key signaling pathways that impacts growth, motility, and survival (6, 13, 21, 22).

Specifically in breast cancer, aberrant HA metabolism is associated with increased tumor invasion and poor patient outcome (23-26). While HA is produced and metabolized by both breast tumor cells and cells in the TME, depleting autocrine HA synthesis in tumor cells results in decreased tumorigenic potential and invasion (27), emphasizing the importance of tumor cell-derived HA in malignant breast cancer progression (26). CD44 and RHAMM are two HA receptors functionally associated with breast cancer progression and metastasis (28, 29). In contrast to CD44, which is ubiquitously expressed (30), RHAMM (gene *HMMR*) expression is low under homeostatic conditions and is upregulated in response to tissue pathologies associated with wound repair, inflammation and/or tumor formation (31-34). Elevated RHAMM expression is associated with higher stage or poor outcome in several cancers, notably breast cancer (35-43).

RHAMM was originally identified as a fibroblast motogenic protein (34, 44), but has been further characterized as a multifunctional oncogenic protein that integrates a multitude of signaling networks to impact tumor cell metabolism, mitosis, cell motility, and invasion (45). RHAMM is an intracellular protein that is unconventionally exported to the cell surface under stress conditions where it interacts with other co-receptors to regulate growth factor receptor (e.g. EGFR, PDGFR) and CD44 signaling (46). RHAMM-CD44 interaction(s) at the cell surface result in oncogenic signaling activated by low molecular weight HA (46-51). When RHAMM remains intracellular, RHAMM is found in both the cytoplasm and nucleus. Cytoplasmic RHAMM associates with the microtubule and vimentin cytoskeleton networks to form dynein motor complexes that localize to the centrosome, which maintains spindle integrity and promotes cellular proliferation and motility (47, 52-54). Nuclear RHAMM forms transcriptional complexes with E2F1 that regulate the expression of genes such as fibronectin (55). While the association of RHAMM expression with diseases such as cancer and its subcellular compartmentalization are well documented, the precise pro-tumorigenic role(s) and mechanism(s) by which RHAMM promotes cancer progression remain poorly understood.

In the present study, we are the first to characterize gene expression signatures that reveal the impact of RHAMM in primary invasive expansion and lymph node metastasis of human breast tumors. We integrate differential gene expression patterns from lymph node positive breast cancer patients with cell models of RHAMM-driven transformation to develop a novel RHAMM Dependent Signature (RDS). We demonstrate that RDS associates with poor outcome in an independent clinical cohort and reveal potential RHAMM-related pathways linked to poor survival. We show that RHAMM protein expression in breast tumors is heterogeneous with distinct regions of strongly positive cells. These RHAMM-high foci are frequently observed near the invasive margin, and digital spatial transcriptomic analysis indicates RDS is also enriched in RHAMM-high regions. Our data show a requirement for RHAMM expression in anchorage independent growth and invasion *in vitro* and invasive progression of *in vivo* intraductal xenografts using CRISPR/Cas9-generated deletion of RHAMM in breast tumor cell lines. We propose that high RHAMM expression in breast tumor cell subsets defines an ‘invasive niche’ in breast cancer that is targetable and will enable development of therapies to limit malignant progression and improve patient outcomes.

## RESULTS

### RHAMM expression is correlated with clinicopathological features of aggressive breast cancers

We explored transcriptional regulation of pathways related to tumor microenvironment (TME) interactions, immune milieu, hormone and growth factor oncogenic signaling, and PAM50 gene expression in a retrospective patient cohort of human breast cancer (n=94) from the University of Minnesota (UMN) Medical Center using a custom Nanostring gene expression probe set (Supplemental Data File 1). We found that increased RHAMM transcript level is significantly associated with lymph node (LN) metastasis (Figure 1A), higher Nottingham tumor grade (Figure 1B), and more aggressive breast cancer subtypes by both PAM50 molecular classification (Figure 1C) and clinical hormone receptor (HR) expression (Figure 1D). This result is consistent with a role for RHAMM function in breast tumor progression.

**Figure 1.**
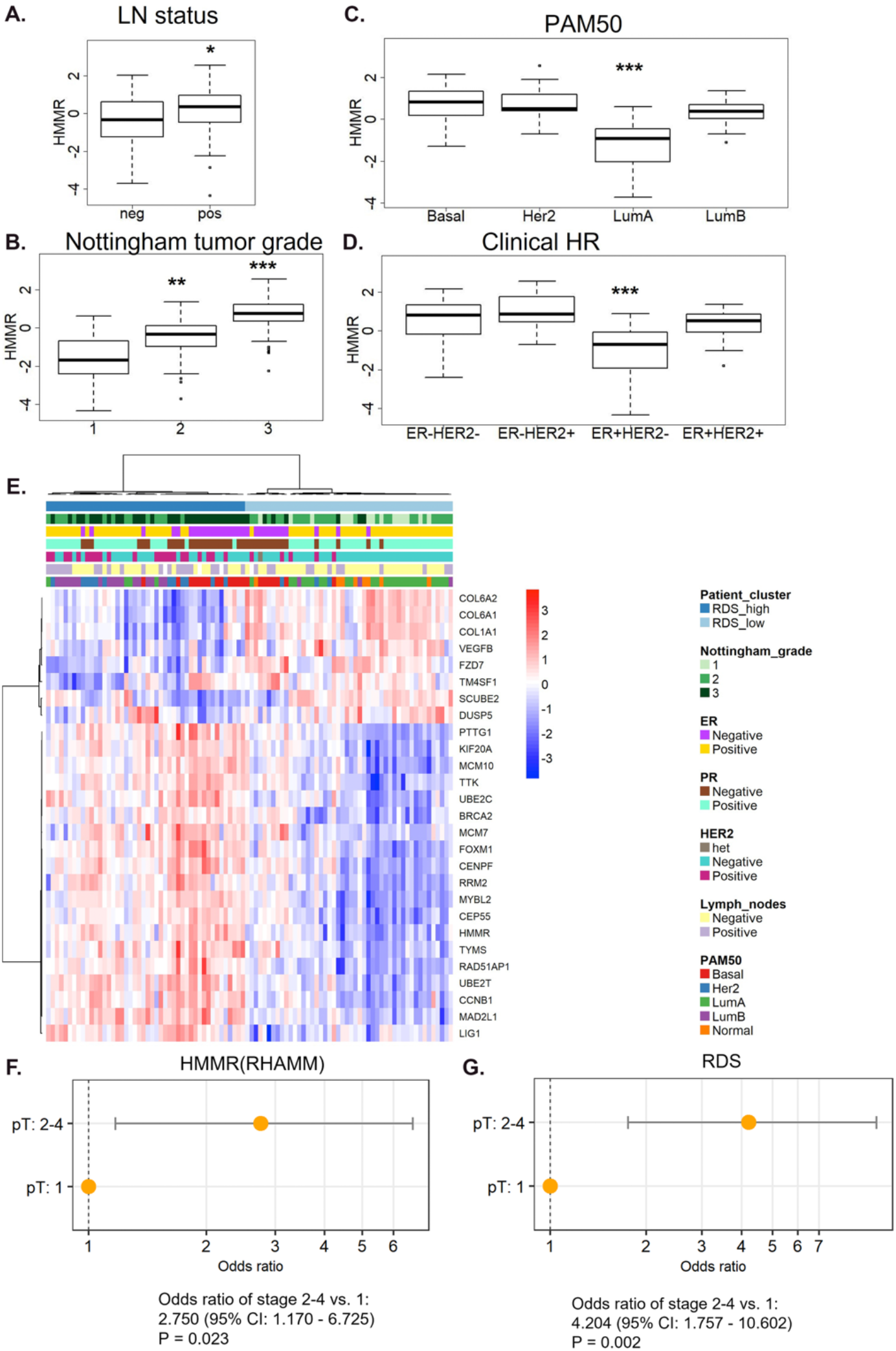
RHAMM expression is correlated with clinicopathological features of aggressive breast cancers. Human breast cancer tissues from the UMN breast cancer cohort were analyzed by Nanostring gene expression. Levels of HMMR mRNA were increased in association with: (**A**) lymph node positive status, (**B**) increased Nottingham tumor grade, (**C**) basal, HER2-enriched, and Luminal B subtypes by PAM50, and (**D**) triple negative and HER2+ disease by clinical hormone receptor expression. (**E**) Unsupervised hierarchical clustering of 94 human breast cancer cases with a 27-gene signature of RHAMM biologic activity. (**F**) Logistic regression analysis of T-stage of RHAMM high vs low patients. (**G**) Logistic regression analysis of T-stage of RDS high vs low patients. * = p<0.05, ** = p<0.01, *** = p<0.005.

To better understand the molecular mechanism(s) of RHAMM in tumor progression we identified the top 50 differentially expressed genes between LN+ versus LN-patients from the UMN patient cohort. We also utilized differentially expressed genes between Rhamm-transformed 10T1/2 and benign parental controls, where overexpression of an oncogenic Rhamm isoform increased cell motility, *in vivo* tumor engraftment, and metastasis (56) to further specifically identify oncogenic RHAMM-related transcripts. Intersection of these two gene sets resulted in a 27-gene RHAMM-dependent signature (RDS) (Supplemental Table 1), which encompasses potential biological consequences of RHAMM activation and signaling in primary human breast cancers with metastatic potential. Pathway enrichment analysis (57, 58) of the RDS showed significant association with cell cycle-related pathways including: E2F1 targets, G2-M checkpoint, mitotic spindle regulation, and MYC targets. Also enriched were ECM-receptor interactions, focal adhesion, and epithelial-mesenchymal transition pathways (Supplemental Table 2). Hierarchical clustering analysis of the UMN patient cohort with RDS genes resulted in RDS-high and RDS-low subsets (Figure 1E). Like RHAMM univariate expression, the RDS-high cluster was significantly enriched for cases with higher Nottingham tumor grade (Supplemental Figure 1A), lymph node metastasis (Supplemental Figure 1B), and more aggressive PAM50 subtypes (Supplemental Figure 1C). We then assessed the probability of either elevated RHAMM expression or RDS-high status to associate with larger invasive tumor size (as defined by clinical pT stage). We found that patients with larger invasive primary tumors have significantly higher odds to have high RHAMM (Figure 1F) and RDS (Figure 1G), with the RDS signature demonstrating a higher odds ratio (4.204) compared to RHAMM alone (2.750). Overall, these data indicate that increased RHAMM expression associates with regulation of pathways related to proliferation, motility, and ECM interaction that correlate with larger invasive tumor growth and metastasis to regional lymph nodes.

Immunohistochemical (IHC) staining for RHAMM protein in the UMN cohort was then performed to confirm the biologic significance of RHAMM transcript analyses and characterize the spatial distribution of RHAMM expression in the complex TME. Total RHAMM protein expression was greatest in breast tumor cells but varied both by the proportion of positive cells and the stain intensity within individual cells (Figure 2A&B). H-scores (59, 60) were quantified by a pathologist to summarize heterogeneous RHAMM protein expression across full histologic sections. The agreement between transcript levels and H-scores across the UMN cohort was significant (R = 0.66, p = 2.6e-11) validating the utility of mRNA expression to estimate increased total protein expression (Supplemental Figure 2A). Thus, increasing RHAMM H-scores trend with increased LN metastasis (Figure 2C), and are significantly associated with higher Nottingham tumor grade (Figure 2D), and more aggressive breast cancer subtypes by both PAM50 molecular classification (Figure 2E) and clinical HR expression (Figure 2F) similar to RHAMM transcript levels. Importantly, we frequently noted focally enriched RHAMM expression in distinct areas of the invasive tumor margin (Supplemental Figure 2B-G). We therefore performed digital spatial profiling of a representative case of high-grade breast cancer that showed heterogenous RHAMM expression with low RHAMM protein expression in the tumor core (Figure 2 G&H) compared to the tumor margin with high RHAMM protein expression (Figure 2G&I) using the Nanostring Cancer Transcriptome Atlas (CTA). Hierarchical clustering of regions of interest (ROIs) from the core and margin with the RDS (limited to 17 genes in the CTA) demonstrated an enrichment of the RDS gene expression at the tumor margin when compared to ROIs at the core of the same tumor (Figure 2J). These data suggest that focal enrichment of RHAMM+ tumor cells at the tumor margin are linked to pathways associated with RHAMM activation and positively correlate with breast cancer progression to larger invasive tumor sizes and potentially LN metastasis.

**Figure 2.**
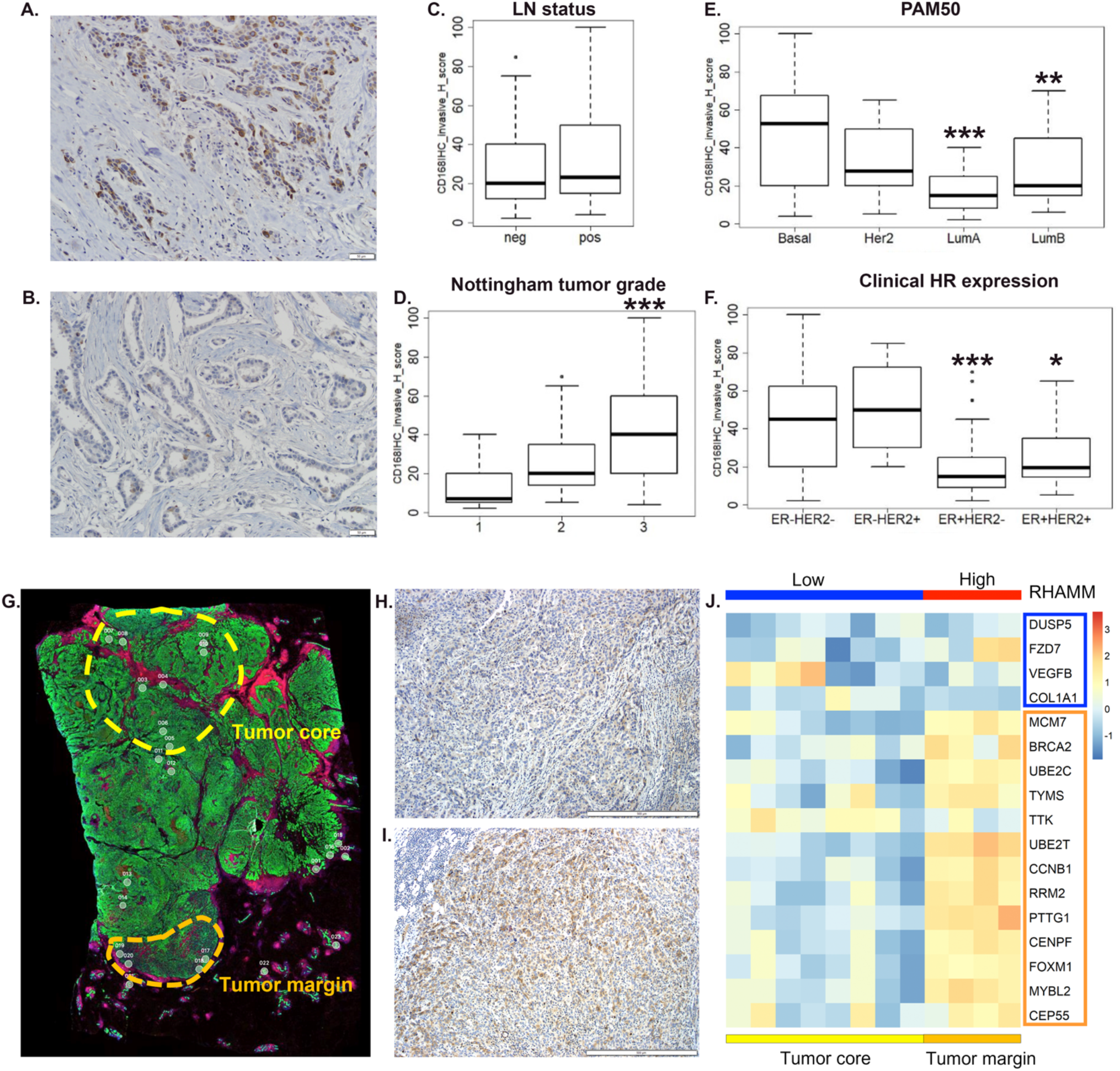
RHAMM protein and RDS expression is heterogeneous across patient samples but correlates with aggressive breast cancers. Representative RHAMM high (**A**) and low (**B**) expressing tissues. (**C-F)** RHAMM IHC were analyzed with high RHAMM levels correlating with: (**C**) lymph node positive status, (**D**) increased Nottingham tumor grade, (**E**) basal, HER2-enriched, and Luminal B subtypes by PAM50, and (**F**) triple negative and HER2+ disease by clinical hormone receptor expression. (**G**) Overview image of a TNBC tumor stained for pan-cytokeratin (green), CD45 (red), and dsDNA (blue). Regions of interest (ROIs) are numbered 1-23 with those in the tumor margin or tumor core highlighted. Serial sections of tumor in panel (**G**) stained for RHAMM at the tumor core (**H**) or the tumor margin (**I**). (**J**) Hierarchical clustering of modified RDS using ROIs located at the tumor core or tumor margin. * = p<0.05, ** = p<0.01, *** = p<0.005,

### RHAMM drives breast cancer invasion

To directly identify how RHAMM contributes to invasive breast cancer progression, we generated RHAMM KO cell lines using CRISPR/Cas9 technology in MCF10DCIS.com cells, a transformed, hormone-receptor negative mammary epithelial cell line which is an ideal model for studying breast cancer progression to invasive disease (61). Immunoblot analysis confirm complete loss of RHAMM protein expression in two independent KO lines (Figure 3A). Although RHAMM can associate with the ubiquitously expressed HA receptor CD44, RT-PCR and flow cytometry analysis shows that CD44 expression is not significantly altered by RHAMM-loss (Figure 3B, Supplemental Figure 3A). Immunofluorescence staining comparing permeabilized and unpermeabilized cells demonstrates RHAMM expression at the cell membrane, within the cytoplasm, and on mitotic spindles in parental and control cells, and confirms complete loss of RHAMM expression in KO cells (Figure 3C). Loss of RHAMM did not significantly affect tumor cell production of HA nor alter HA fragmentation (Supplemental Figure 3B&C). Further, cell internalization of exogenous low molecular weight (LMW) HA added to the culture medium was not impacted by RHAMM loss (Supplemental Figure 3D&E). Collectively, these data demonstrate specific stable knockout of RHAMM without significant perturbation of CD44 expression or HA metabolism *in vitro*. We next assessed the effects of RHAMM KO on breast cancer progression phenotypes *in vitro*. Proliferation was not significantly impacted by RHAMM KO (Figure 4A). Conversely, RHAMM loss significantly decreases tumor cell ability to grow in anchorage independent conditions (Figure 4B). Further, RHAMM KO significantly decreases invasion of breast cancer cells in a 3D invasion assay (Figure 4C). Finally, vimentin protein expression, a biomarker associated with mesenchymal plasticity and motility (62), was consistently decreased in RHAMM KO cells (Figure 4D). Taken together, these data suggest RHAMM increases breast tumor cell invasion and anchorage independent growth.

**Figure 3.**
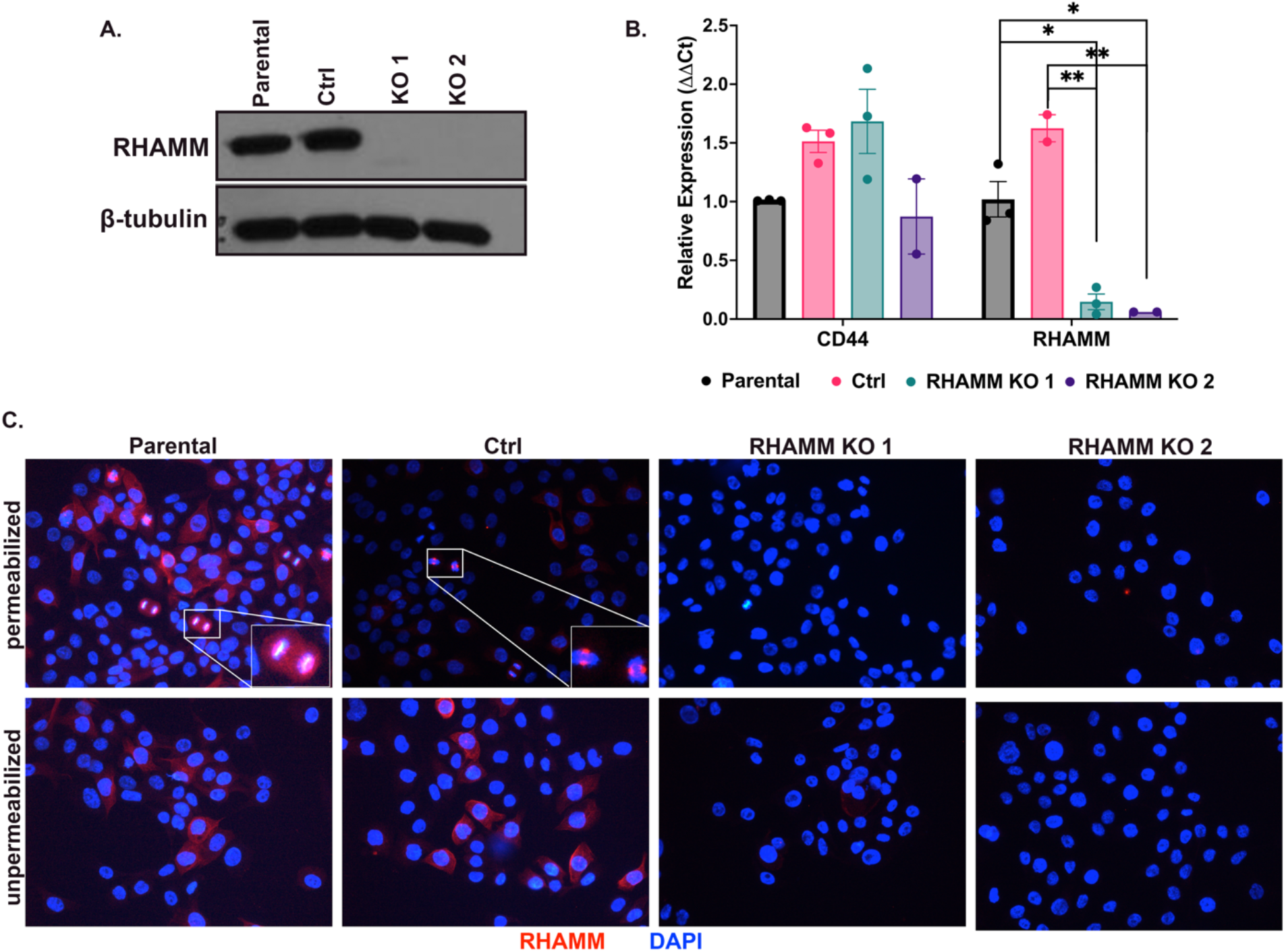
Characterization of RHAMM KO breast cancer cells. (**A**) Representative immunoblot of MCF10DCIS.com parental, Ctrl, and RHAMM KO cell lines for RHAMM. (**B**) Quantitative-PCR expression of CD44 and RHAMM in MCF10DCIS.com parental, control and RHAMM KO cell lines. (**C**) Representative RHAMM immunofluorescence images of MCF10DCIS.com parental, control and RHAMM KO cell lines, in permeabilized and unpermeabilized conditions, insets highlighting mitotic spindle RHAMM staining. * = p<0.05, ** = p<0.01.

**Figure 4.**
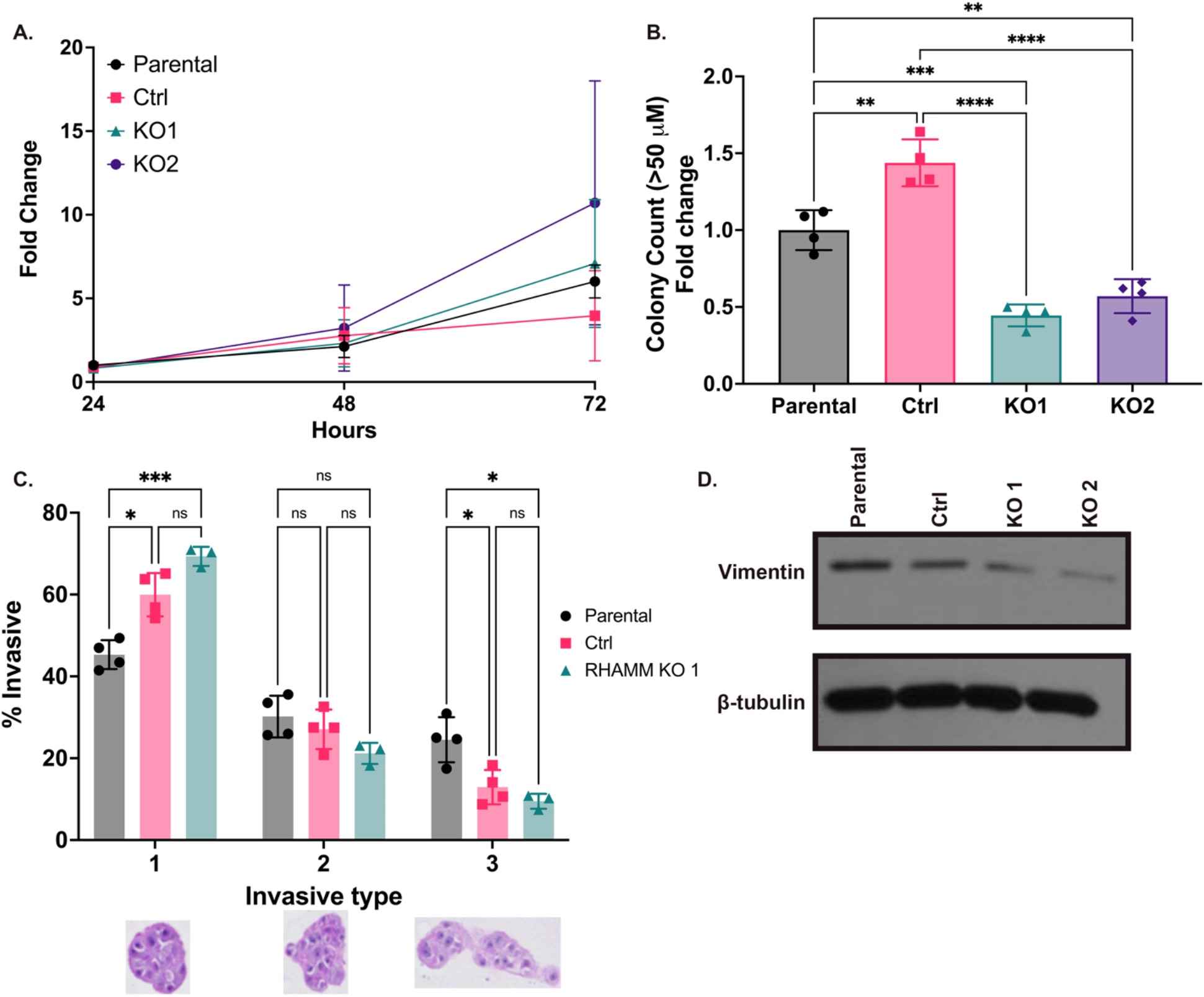
RHAMM drives phenotypes associated with invasive breast cancer. (**A**) MTT proliferation assay of MCF10DCIS.com parental, control and RHAMM KO cell lines. (**B**) Colony counts (size >50μm) of MCF10DCIS.com parental, control and RHAMM KO cell lines in soft agar. (**C**) MCF10DCIS.com parental, control and RHAMM KO cell lines embedded in Matrigel + 20% collagen and scored for invasion, representative images of tumorspheres below. (**D**) Representative immunoblot of vimentin in MCF10DCIS.com parental, control and RHAMM KO cell lines. * = p<0.05, ** = p<0.01, *** = p<0.005, **** = p<0.001

To directly assess this in a complex TME, we next performed *in vivo* studies using the mouse mammary intraductal (MIND) model where, tumor xenografts undergo a period of intraductal proliferation, mimicking progression of human ductal carcinoma *in situ* to invasive breast cancer (61, 63). We injected parental, control, and RHAMM KO1 MCF10DCIS.com cells into the ducts of the fourth mammary gland of NSG mice. After approximately 4 weeks, wildtype xenografts undergo reliable invasive progression and expansion into palpable tumors (Supplemental Figure 4A) while retaining variable amounts of residual *in situ* tumor foci. At an endpoint of six weeks, the majority of parental and control tumors reached maximum allowable tumor volume (2000 mm^3^) while all KO xenografted animals survived with minimal to no grossly appreciable tumor burden (Figure 5A). Subsequent histologic analysis reveals that parental and mock control tumors showed almost complete progression to invasive ductal carcinoma (IDC), with 75% of parental and 92.9% of control tumors progressing to predominantly IDC (Figure 5B). Conversely, 0% of RHAMM KO tumors progressed to IDC. The majority, 50%, of RHAMM KO tumors never progressed beyond *in situ* compared only 16.7% and 0% in parental and control tumors, respectively. RHAMM KO tumors that progressed to invasive growth had significantly smaller maximum invasive tumor sizes (Figure 5C) and correspondingly lower gross tumor volume volumes and weights (Supplemental Figure 4B&C). Further IHC analysis reveals a significant increase in the proportion of RHAMM-positive tumor cells in the areas of invasive growth when compared to *in situ* areas in parental (RHAMM expressing) tumors (Figure 5D). This observation is consistent with an upregulation of RHAMM supporting invasive expansion during breast cancer progression. Quantification of the Ki67 labelling index showed no significant differences in the percentage of Ki67+ cells between parental, control, and KO tumors in either *in situ* (Supplemental Figure 4D) or invasive tumors (Supplemental Figure 4E), indicating that genomic *RHAMM* deletion did not cause a quiescent tumor cell phenotype *in vivo*. However, assessment of the mitotic-phase marker phosphorylated histone H3 (pH3) (64-67) show significant differences in the average number of pH3+ cells per 10 high power fields in the invasive areas of *RHAMM* wildtype controls compared to invasive KO xenografts (Supplemental Figure 4F), indicating that RHAMM may promote more frequent mitotic cell division *in vivo*. Hyaluronan, measured by HAPB, was noted to accumulate at the leading edge of invasive tumors, variably within the tumor core, and at the periphery of DCIS lesions (Figure 5E, Supplemental Figure 4G&H). Interestingly, we observed higher intra-tumoral HA accumulation in invasive components of RHAMM KO tumors (Figure 5E), which may suggest more complex alterations in HA metabolism with loss of RHAMM *in vivo*, not detected *in vitro*. Overall, these data demonstrate that RHAMM expression is required for breast tumor invasion.

**Figure 5.**
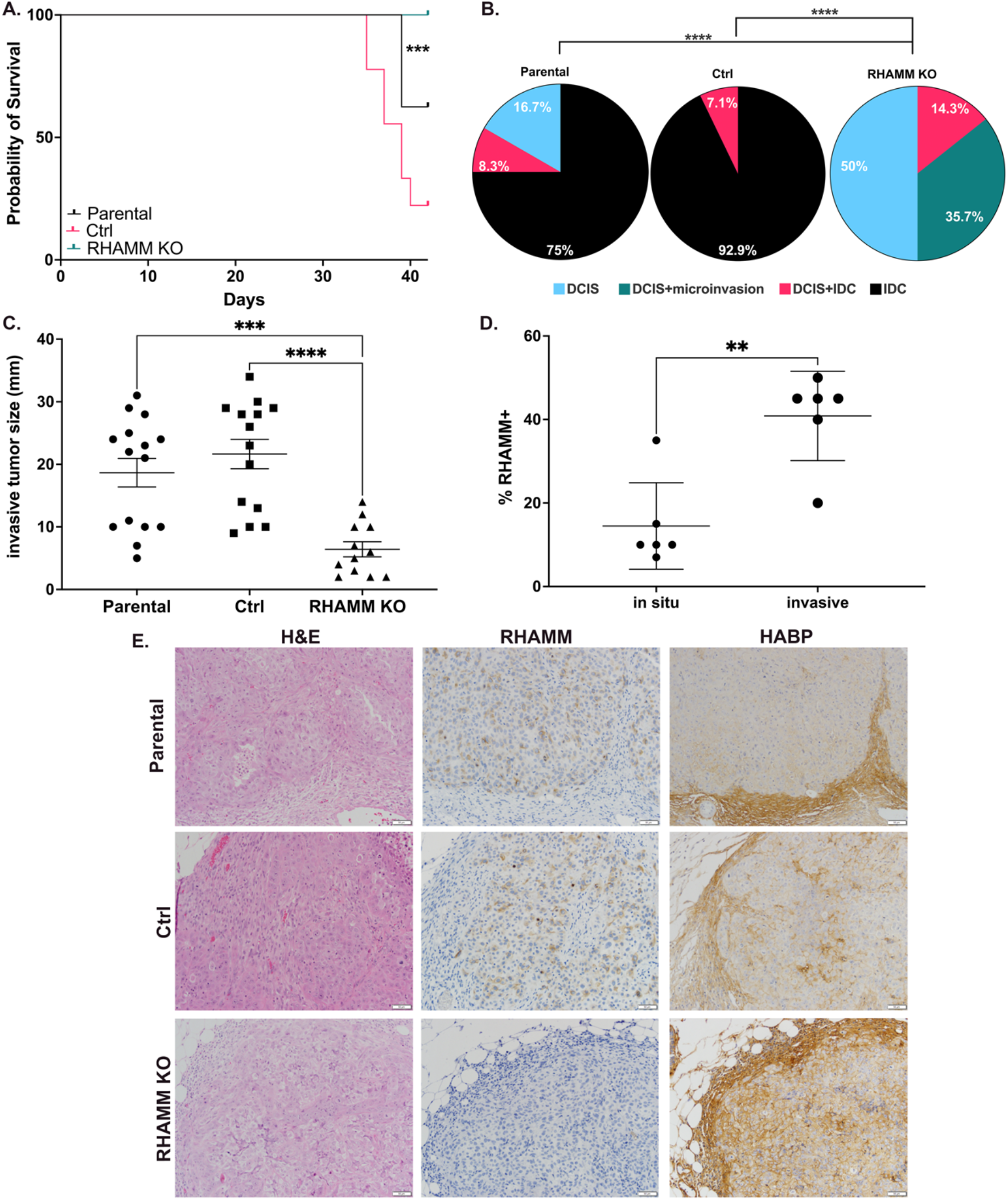
RHAMM drives breast tumor invasion. (**A**) Survival analysis of mice bearing tumors from MCF10DCIS.com parental, control and RHAMM KO cell lines, with survival calculated by time to tumor size of 1000 mm^3^. (**B**) Tumors from MCF10DCIS.com parental, control and RHAMM KO groups scored for invasion. (**C**) Size of invasive tumors from parental, control, and RHAMM KO cell lines. (**D**) MCF10DCIS.com parental tumors scored for invasion and then stained by IHC and scored for RHAMM expression. (**E**) Representative images of invasive tumors stained for H&E and IHC for RHAMM and HABP. ** = p<0.01, *** = p<0.005, **** = p<0.001.

### RHAMM Dependent Signature (RDS) correlates with poor outcomes in breast cancer patients

To further validate the clinical significance of our findings we analyzed microarray expression data from the Breast Cancer Collaborative Registry (BCCR) at the University of Nebraska. Corroborating our previous results with the UMN dataset (Figure 1E), unsupervised hierarchical clustering of the BCCR dataset also identifies a subgroup of RDS high patients who are enriched for aggressive clinicopathological features (Figure 6A, Supplemental Figure 5 A-C). Importantly, RDS-high patients have worse 10-year overall survival than RDS-low patients in the BCCR dataset (Figure 6B). The integration of RHAMM and downstream/associated pathways in the RDS better identifies this prognostic phenotype than measurement of *RHAMM* transcript expression alone (Supplemental Figure 5D). Additional analysis using the TCGA Breast Cancer dataset also reveals that RDS predicts for worse overall survival (Figure 6C). Overall, our findings define RHAMM as a key mediator of breast cancer progression and we propose a novel gene expression signature, RDS, may identify aggressive breast cancers with increased capacity for larger invasive primary tumor expansion, metastatic progression, and ultimately worse overall survival. Identification of RDS-enriched breast cancers will enable further study of mechanisms by which RHAMM and HA activate invasive progression.

**Figure 6.**
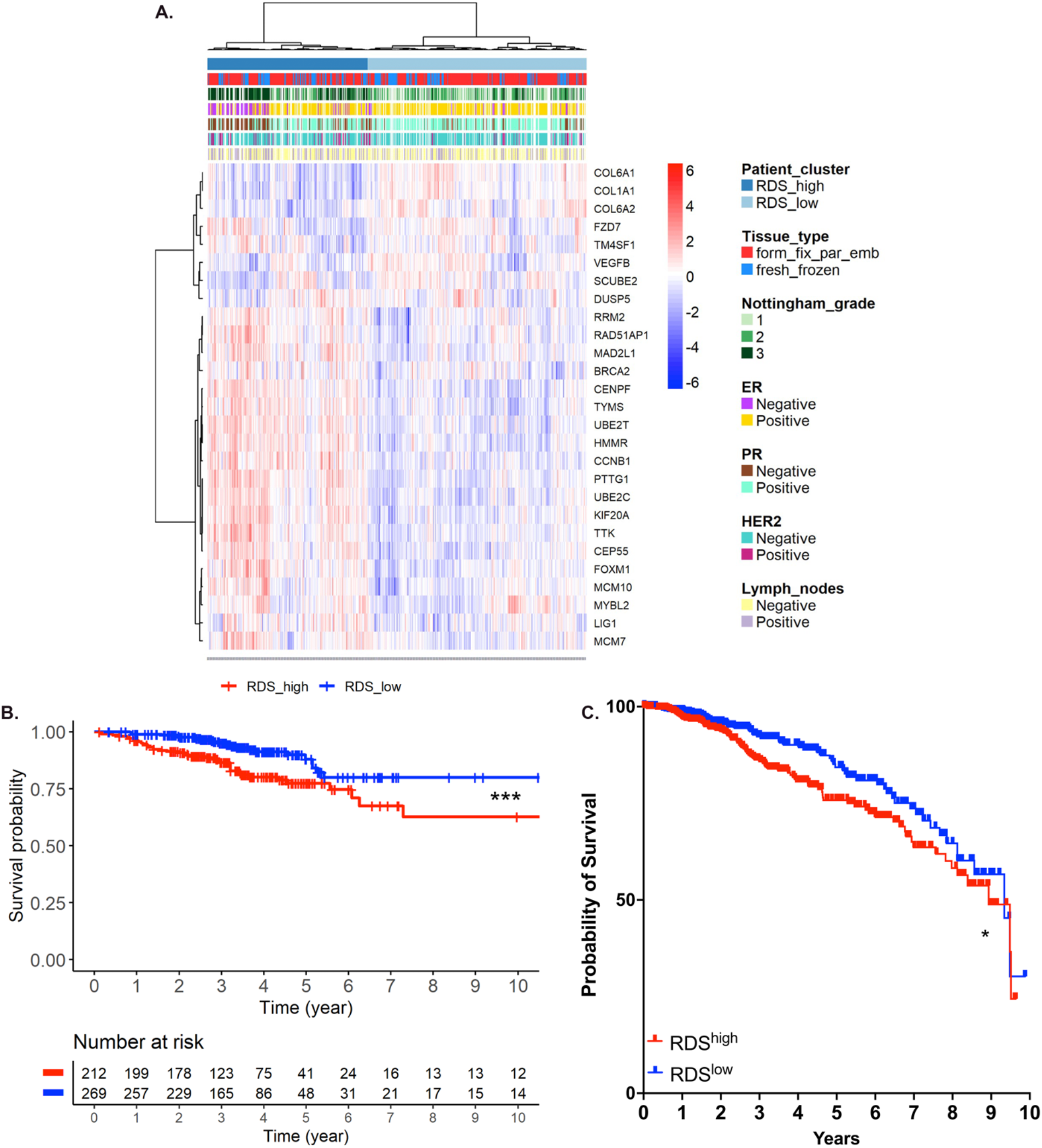
RDS correlates with poor outcomes in breast cancer patients. (**A**) Unsupervised hierarchical clustering of human breast cancer cases from the University of Nebraska Breast Cancer Collaborative Registry (BCCR) with a 27-gene signature of RHAMM biologic activity (n=656). (**B**) Kaplan-Meier plot of BCCR cases by RDS high or low for overall survival at 10 years (n=481). (**C**) Kaplan-Meier plot of TCGA breast cancer cases by RDS high or low for overall survival at 10 years (n=1035).

## DISCUSSION

Accumulating evidence suggests that interactions between tumor cells and their cancerized microenvironment are critical for understanding mechanisms driving cancer progression, which are essential for developing new therapeutic interventions. Herein we use a discovery breast cancer patient cohort that verifies the association of *RHAMM* transcript with aggressive clinicopathologic features and identify potential downstream effector pathways of RHAMM activity. RHAMM is a hyaluronan receptor that is dynamically regulated in contexts of cell stress which enables cells to sense and respond to cues in the extracellular matrix (11, 68). Elevated *RHAMM* expression was significantly associated with lymph node positivity, high Nottingham tumor grade, and more aggressive molecular and clinical subtypes of breast cancer. We propose a novel RHAMM Dependent Signature (RDS) informed by differentially expressed genes in primary breast tumors with lymph node metastasis intersected against the transcriptome of RHAMM-dependent cell transformation. Analysis of independent breast cancer patient cohorts validates that RDS associates with worse overall breast cancer survival. RDS-high patients additionally have larger invasive tumor size and is enriched across more aggressive subtypes including triple negative breast cancer TNBC/basal, HER2+, and Luminal B, suggesting that RDS encompasses common pathways used by diverse breast cancer subtypes to drive progression.

Immunohistochemical analysis of tumors in the UMN cohort showed that RHAMM protein expression is heterogeneous within the breast tumor microenvironment, with only subsets of tumor cells demonstrating moderate-high RHAMM staining levels, frequently at the invasive tumor margin. Digital spatial transcript profiling of human TNBC indicates that niches of RHAMM+ tumor cells at the invasive tumor margin express a RDS-high gene profile compared to RHAMM-low areas within the same tumor, showing that this pattern of gene expression is directly linked to focal regions of increased RHAMM protein expression. These spatially-resolved observations from human breast cancer support our hypothesis that micro-anatomically distinct foci of RHAMM+ tumor cells form localized niches with unique downstream pathway activation that drives tumor cell invasion.

The capacity for invasion and motility is the first and irrevocable hallmark of metastasis (3). After losing contact inhibition and anchorage dependency, tumor cells’ ability to erode and escape through the basement membrane is the defining characteristic for classification of invasive malignancy (3). In this study, genomic knockout of *RHAMM* in a model of breast cancer progression decreases tumorsphere invasion and anchorage independent growth *in vitro*, and dramatically decreases xenograft tumor invasive progression and expansion *in vivo*. A limitation of the MCF10DCIS.com xenograft model is that it does not generally metastasize well.

Additional studies will need to address the role of RHAMM and how its activation may contribute to distant metastasis. Since we observe significant enrichment of RDS and elevated *RHAMM* in clinical LN+ disease, we suspect RHAMM and downstream signaling will play a role in multiple steps of the metastatic cascade. In the parental and control xenografts we observe RHAMM expression increases endogenously during the progression from *in situ* to invasive growth along with simultaneous HA accumulation around DCIS lesions and at the invasive tumor margin. Alone, HA accumulation has been associated with breast cancer progression, and pharmacological inhibition of HA synthesis has been shown to enhance apoptosis or have other anti-tumor effects in solid tumor models (6, 69-71). These observations are consistent with a role for RHAMM-HA interactions in promoting invasive expansion. Of note, RHAMM KO cells maintain *CD44* expression, suggesting RHAMM may act in a unique manner distinct of other HA-sensing molecules and HA-machinery to promote breast cancer progression. Further, since RHAMM lost does not alter HA internalization CD44 or other HA receptors may be acting in a compensatory mechanism. Taken together our studies suggest that targeting RHAMM function may have therapeutic benefit for breast cancer patients.

The development of mesenchymal characteristics, decreased cell-cell adhesion, and stem-like properties is described as epithelial-mesenchymal transition (EMT), an important contributing, but not necessarily required, phenotype for invasive progression and metastasis (3). EMT and its converse mesenchymal-epithelial transition are not necessarily distinct binary states, but rather tumor cells likely exist on a continuum between these two poles with dynamic expression of both epithelial and mesenchymal markers (3). This phenotypic plasticity is an emerging hallmark of cancer that supports malignant progression (72). In our model, RHAMM KO decreases vimentin expression and limits invasive or anchorage independent growth without significantly impairing proliferation *in vitro*. RHAMM KO xenograft tumors do not have significantly decreased Ki67 in either *in situ* or invasive components, suggesting that the impaired rates of progression and invasive expansion are not due to increased cell quiescence. In comparison, the mitotic marker pH3 was significantly higher in wildtype tumors, suggesting that RHAMM wildtype xenografts were more frequently undergoing mitotic division *in vivo*, consistent with the larger observed invasive tumor sizes. The increased number of cells in G2/M in wild type and tumors may also increase *in vivo* motility (73), potentially mediated by intracellular RHAMM interactions with cell cycle machinery. In this context, we propose that RHAMM may promote tumor cell plasticity, which allows for more rapid invasive tumor expansion.

Along these lines, RDS encompasses pathways of cell cycle regulation/E2F/MYC activity, as well as EMT, ECM-receptor interaction, and cell adhesion. The utilization of an oncogenic RHAMM transcriptome from a mesenchymal cell model may have highlighted potential pathways of mesenchymal phenotypic plasticity when developing this gene signature. Further, the E2F pathway and other mitotic regulators are known to have roles in the regulation of EMT (74, 75), and RHAMM-E2F1 association may modulate the microenvironment through upregulation of fibronectin to support the metastatic cascade (55). These observations lend further support to a model that RHAMM supports phenotypic plasticity to enable invasive progression through coordination of both intracellular and extracellular oncogenic mechanisms.

We propose that RHAMM protein upregulation defines distinct niches within breast tumors that serve as drivers of invasive progression. Breast tumors with a greater density of active RHAMM+ invasive niches have a greater capacity for both local invasive tumor expansion and lymph node metastasis. Future studies need to address how RHAMM is upregulated in these niches and mechanistically dissect which downstream activities are necessary to support an invasive phenotype. Further work should characterize supporting stromal cells in the RHAMM+ niche, including the phenotypes of cancer-associated fibroblasts and tumor-associated macrophages therein. Our results suggest that targeting RHAMM function or RHAMM:HA interaction may serve as an additional therapeutic mechanism to attack tumor growth and limit metastatic spread. This is important because RHAMM/RDS upregulation is common in several molecular subtypes of aggressive breast cancers, including TNBC, this study demonstrates a potential target to complement existing standards of care.

## MATERIALS AND METHODS

### Cell Culture

MCF10DCIS.COM cell lines (provided by Dr. Fariba Behbod, University of Kansas Medical Center) were sub-cultured in Dulbecco’s Modified Medium/Ham’s Nutrient Mixture F-12 (Thermo Fisher Scientific, Cat.# 51448C) base media containing 5% Horse Serum with 1% Penicillin/Streptomycin at 37ºC and 5% CO_2_. Cells were regularly tested for mycoplasma throughout studies and validated against ATCC STR profiles. RHAMM KO cell lines were generated using CRISPR/Cas9 technology, with gDNA plasmids obtained from Dr. Branden Moriarity. Following transfection, cells were clonally selected and tested for RHAMM KO by DNA and protein. 3D culture assays were performed as previously described (76, 77).

### MTT Proliferation Assay

Cells were plated in a 96-well plate at 2000 cells/well in 200ul complete growth medium. For each timepoint, MTT reagent (5mg/mL, Thermo Fisher Scientific, Cat.#: M6494, MA, USA) was added to each well and incubated for 2 hours at 37ºC protected from light. Solubilization buffer (95% DMSO in PBS) for 10 minutes followed by reading absorbance at 540 nm.

### Immunofluorescence

Cells were cultured in an 8 well chamber slide (Sigma Cat.#: C7057) then fixed in 4% paraformaldehyde and where appropriate permeabilized with 2% TritonX-100. Cells were then stained with RHAMM antibody (Supplemental Table 3), overnight at 4 °C, then appropriate secondary (AF 594 anti-Rabbit) for 1 hour at room temperature. Slides were coverslipped with ProLong Gold Antifade DAPI (Invitrogen, #P36931).

### Microscope Imaging

All images were acquired on a Leica DM400B microscope (Leica, Wetzlar, Germany), at either 20× or 40× objectives. Images were acquired using a Leica DFC310 FX camera (Leica, Wetzlar, Germany) and LAS V3.8 software. Five images of at least 3 representative tumors were analyzed.

### HA Fragmentation Analysis

Analysis of HA fragmentation via polyacrylamide gel electrophoresis was performed as previously described (78) using concentrated cell culture conditioned medium.

### Anchorage Independent Colony Formation Assay

Cells were cultured in 0.3% agarose solution for 21 days. Colonies (> 50 µM) were counted under an inverted light microscope.

### Animal Model

8-10 week old, female, NOD-*scid* IL2Rgamma^null^ mice from Jackson Laboratories were injected intraductally (MIND model) as previously described (61). Briefly, following nipple resection, 10,000 cells were injected intraductally bi-laterally into the number 4 mammary gland. Following tumor cell injections, tumors were measured bi-weekly.

### Immunoblotting

Cells were lysed in RIPA buffer with protease inhibitors (Roche: Cat#11836170001). Following complete lysis and lysate clearing, protein was quantified using a DC protein assay (BioRad: Cat#: 5000111). Equal amounts of protein were then loaded on an 8% SDS gel, run, and transferred to a PVDF membrane for immunoblotting. Antibody information is provided in Supplemental Table 3.

### qPCR

RNA was harvested using TriPure Isolation reagent (Sigma: Cat#: 11667165001). Following RNA quantification using a spectrophotometer at 260/280, cDNA was made using qScript cDNA synthesis kit (QuantaBio: Cat #101414-098). qPCR was performed using SYBER-green (VWR: Cat #101414-144) on a BioRad iQ5 Thermocycler. CD44 primers: Forward: 5’ - GAT GGA GAA AGC TCT GAG CAT C -3’ Reverse: 5’ - TTG CTC CAC AGA TGG AGT TG -3’ RHAMM PrimePCR primers were obtained from BioRad (Cat#: 10025636)

### Histology and Immunohistochemistry (IHC)

Breast cancer xenografts were collected, fixed with 10% NBF, and embedded in paraffin. Two H&E stained levels were analyzed by a board certified pathologist (ACN) blinded to genotype for scoring of the proportion of *in situ* (DCIS), microinvasive, and invasive carcinoma (IDC) growth patterns within each tumor section as previously described (76, 79). A standard IHC was performed to evaluate the expression of RHAMM, Ki67, and phospho-histone 3 (pH3). Slides were gapped with a 6.0 pH citrate buffer for heat retrieval using a vegetable steamer for 30 min and cooled down at room temperature for 20 min. Then the slides were loaded onto the Intellipath IHC platform for staining. For HABP IHC slides were rehydrated using standard methods. Slides were then placed into TBST buffer.

Following buffer, the slides are divided into 2 sets, one set is incubated with a 1% hyaluronidase and one set with TBST at 37°C for 30 minutes. Slides were then rinsed in TBST and blocked in Sniper blocking (Biocare Medical) one hour at room tmperature. Anti-bHABP (Calbiochem) was applied to all slides and incubated overnight at 4°C. The next day slides were rinsed and avidin-biotin-complex (ABC, Vector Labs) was applied for one hour at room temperature. Slides were rinsed and counterstained with CATHematoxylin diluted 1:2 (Biocare Medical) for 5 minutes, rinsed, dehydrated and cover slipped. Antibody information provided in Supplemental Table 3.

### Flow cytometry

Cells were harvested via scraping. Cells were then filtered through a 40 μM filter and blocked via FcR (FACS) buffer. Cells were stained with using Zombie NIR™ fixable viability kit and CD44 (Supplemental Table 3). Flow cytometry was performed on Cytek Aurora machine. All flow cytometric events defined by FSC x SSC forward and side scatter, doublet discrimination, and viable cells defined by exclusion of viability dye. Cells were then gated on minus debris and singlets. Data analysis was performed with FlowJo.v10.7.2

### HA binding assay

Cells were harvested using Accutase (Cat # 423201, BioLegend, CA, USA) and filtered through a 40 µm strainer. Preexisting HA matrices were removed via hyaluronidase (2.5 mg/mL, Cat # H3506, MilliporeSigma, MD, USA) treatment for 30 min at 37ºC. Cells were then stained for viability using Zombie NIR™ fixable viability kit for 15 min at room temperature (Cat # BioLegend, CA, USA), followed by fluorescently labeled high- or low-molecular weight HA (2.5mg/ml in PBS) for 1 hour at room temperature. Flow cytometry was performed on a BD LSR Fortessa H0081 machine. Data analysis was performed with FlowJo.v10.7.2.

### HA internalization assay

Cells were seeded in chamber slides (Cat # 80841, IBIDI). 48 hours post-plating, cells were incubated with TexasRed labeled high molecular HA (Cat #H-700R, Echelon Biosciences) or low molecular HA (Cat #H-025R, Echelon Biosciences) in fresh medium for 45 min at 37 °C in dark. Then they were washed with PBS and fixed with 2% PFA for 10 minutes in room temperature in dark. They were subsequently permeabilized by 200ul 2% Triton X-100 in PBS at room temperature for 30 minutes. Then they were stained with 200 ul Phalloidin (1:500 dilution, Cat # 501720934, Thermo Fisher Scientific, MA, USA) for 1 hour at room temperature. After washing and drying, ProLong™ gold antifade mount with DAPI was added to the slides. Images were acquired on a on a Leica DM400B microscope (Leica, Wetzlar, Germany) at 40× objective.

### Digital Spatial Profiling

One formalin-fixed paraffin-embedded (FFPE) 5 µm section of triple-negative breast cancer was obtained from the Department of Lab Medicine and Pathology, University of Minnesota. The informed consent was signed by the patient prior to this study. The slide was prepared using the GeoMx DSP RNA Slide Prep Kit for FFPE (NanoString, Seattle, WA, USA) following the manufacturer’s instructions. In situ hybridization of GeoMx Cancer Transcriptome Atlas (NanoString, Seattle, WA, USA) probe set was performed as directed by the manufacture’s protocol. Then the slide was stained with morphology markers using the GeoMx Morphology Kit for RNA (NanoString, Seattle WA). After wash, the slide was loaded onto the GeoMx DSP instrument to select regions of interest (ROIs) by a pathologist (ACN) in conjunction with a serial section immunohistochemically stained for RHAMM. ROIs were collected and RNA expression profiling according to manufacturer’s’ instructions (NanoString, Seattle, WA, USA). Raw read counts were uploaded to the GeoMx DSP Control Center (Version 2.3.0.268) for quality control (QC), scaling, and normalization according to the manufactures’ manual. QC was done using the default thresholds recommended by the manufacture (NanoString, Seattle, WA, USA). QC flagged ROIs were excluded from the downstream analysis, and QC flagged probes were excluded from all the ROIs. The raw read counts were scaled based on the ratio of the geometric mean ROI area to the measured surface area of each ROI. Then the read counts were further normalized to the 3rd quartile of all selected probes.

### NanoString Gene Expression Analysis

A custom gene set (n=356) was developed covering genes involved in: tumor microenvironment interactions, inflammatory signaling, steroid receptor and growth factor signaling, cell cycle, apoptosis, and DNA repair, plus PAM50 classifier genes. Raw gene expression counts were analyzed within R (ver 3.6.1, R Core Team, 2019) using the NanoStringQCPro package (ver 1.18.0). Flagged outliers (26 cases out of 120 cases) were removed for downstream analysis. After normalization, each gene was median-centered and scaled (root mean square) based on expression levels across all 94 high-quality samples.

### Microarray Gene Expression Analysis

Raw microarray expression data was exported from Feature Extraction software (ver 7.5.1, Agilent Technologies) and analyzed within R using the limma package (ver 3.42.2). Major outliers (n = 20 were removed, leaving 656 samples for downstream analysis. Probes were removed from the dataset if they were not annotated with a gene symbol or were not expressed above background levels in a majority of samples, leaving 24,625 probes included in downstream analyses.

### Gene Expression Plotting

The data were explored by hierarchal clustering of median-centered and scaled gene expression values by calculating a Pearson correlation distance matrix and linked using the Ward1 (ward.D) algorithm in R. Heatmaps were generated using the aheatmap function from the NMF R package (ver 0.23.0). Boxplots were generated by in R that describe the median, hinges correspond to the 1st and 3rd quartiles, and the whisker extends from each hinge 1.5 * IQR.

### Statistics

All in vitro studies were performed in biological triplicates, unless otherwise noted. All animal studies were repeated at least twice with representative data shown. Quantitative experiment values are shown as mean and standard deviations, unless otherwise noted. Survival curves were plotted by Kaplan Meier method and the difference between curves was evaluated by a log-rank test. Differences of the distribution of categorical variables were evaluated by Chi-square test or Fisher’s exact test (when the expected frequency < 5). Differences in quantitative values among experimental groups were evaluated by t test or one way ANOVA (with Tukey’s honestly significant difference post hoc test). Logistic regression was performed to estimate odds ratio and coefficients of categorical dummy variables. P value less than 0.05 is considered statistically significant. All data visualization and statistical analysis were performed using R version 3.6.1 and GraphPad Prism version 8.0.0. TCGA analysis was performed using UCSC Xena Browser (80).

### Study approval

The UMN cohort (IRB Approval Study# 1409E53504) was collected retrospectively from archived pathology tissue blocks with de-identified clinical data in compliance with HIPPA regulations and institutional policies; all participants had agreed to the institution’s standard consent for research participation. The BCCR cohort was collected as previously described (81). All animal studies were approved by the IACUC of the University of Minnesota, protocol number 1810-36470.

## AUTHORS’ CONTRIBUTIONS

ACN, SET, CD, YH, CT, DY, KLS, ET, and JM conceived and designed the research studies. SET, CD, YH, MP, PW, RK, LG, MS, XZ, and CF performed the in vitro and in vivo studies. ACN, CHa, OS, MMD, KC and DY acquired human clinical samples and data and were responsible for regulatory oversight/compliance. TPK, CHe, RS and YH performed bioinformatics and biostatistical analysis. ACN was responsible for analyzing and scoring all tumors for invasion and IHC analysis. SET, CD, YH, and ACN were responsible for hypothesis development, conceptual design, data analysis, and data interpretation. SET and ACN wrote the manuscript with all authors providing critical evaluation. ACN is the guarantor of integrity for the study.

## ACKNOWLEDGEMENTS

ACN is supported by the American Cancer Society (132574-CSDG-18-139-01-CSM). SET is supported by IRACDA-TREM Postdoctoral Fellowship. KS is supported by the NIH R01CA215052, R01HD095858. ET and CT are supported by the Breast Cancer Society of Canada. JBM is funded by the Atwater Fund, Elsa Pardee and the Chairman’s Fund Professor in Cancer Research. This work was supported in part by the Masonic Cancer Center, University of Minnesota and Minnesota Masonic Charities. We acknowledge the Minnesota Supercomputing Institute (MSI) at the University of Minnesota for providing high performance computational resources. This research received histology and immunohistochemistry assistance from the University of Minnesota’s Biorepository and Laboratory Services program and was supported by the National Institutes of Health’s National Center for Advancing Translational Sciences, grant UL1TR002494. The authors wish to thank all the scientists and research support staff in the University of Minnesota Genomics Center, University Imaging Center, and Laboratory Medicine and Pathology who helped to make this work possible.

