## Supplemental Figures for "Receptor for Hyaluronan-Mediated Motility (RHAMM) defines an invasive niche associated with tumor progression and predicts poor outcomes in breast cancer patients"

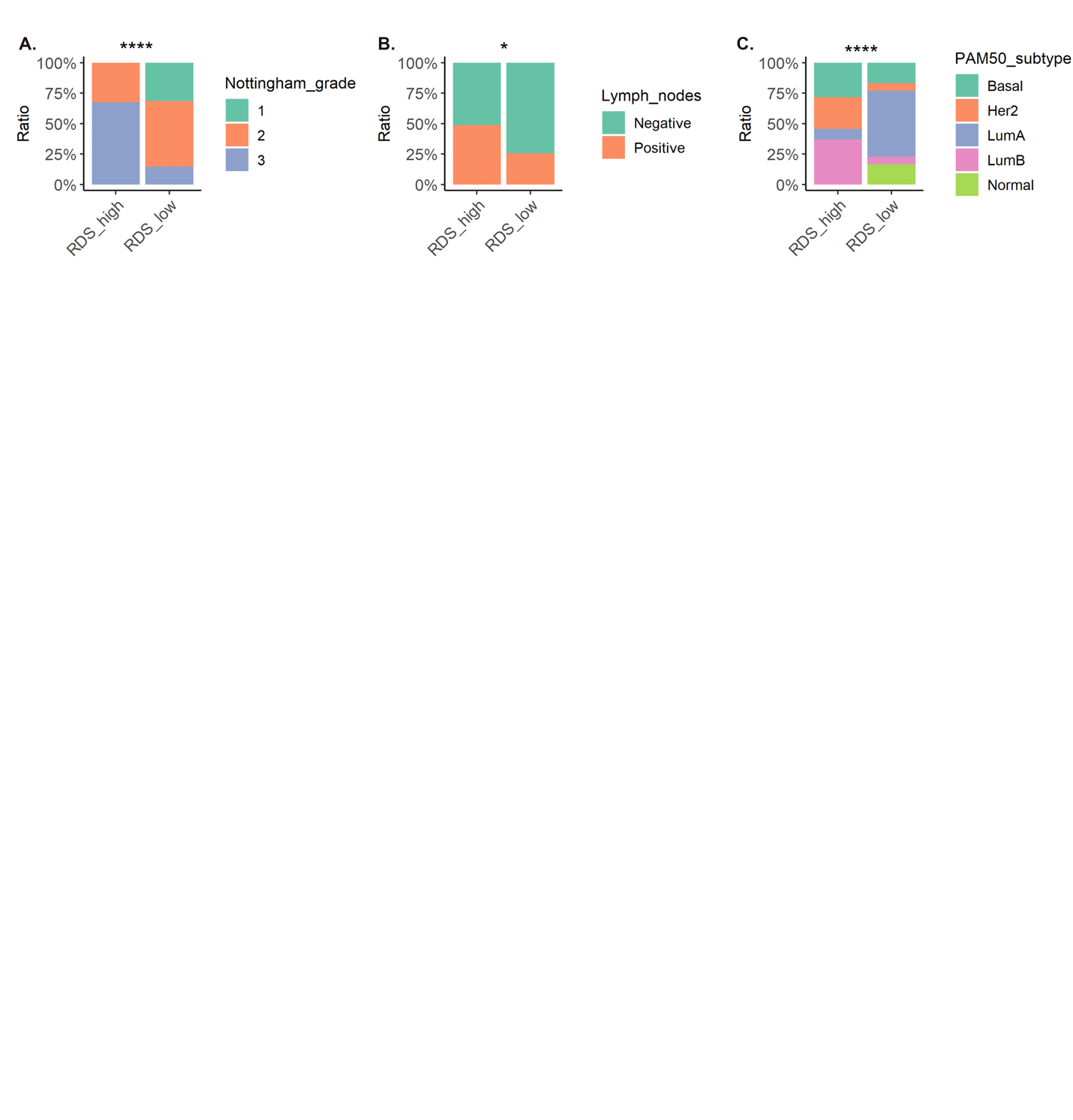

**Supplemental Figure 1**. RDS is correlated with clinicopathological features of aggressive breast cancers. Distribution of clinicopathological features across RDS clusters of breast cancer patients (the UMN datasets), Nottingham grade (A), lymph node status (B), and PAM50 subtype (C). * p value < 0.05; **** p value < 0.0001.

Supplemental Table 1. Genes in RHAMM Dependent Signature (RDS)

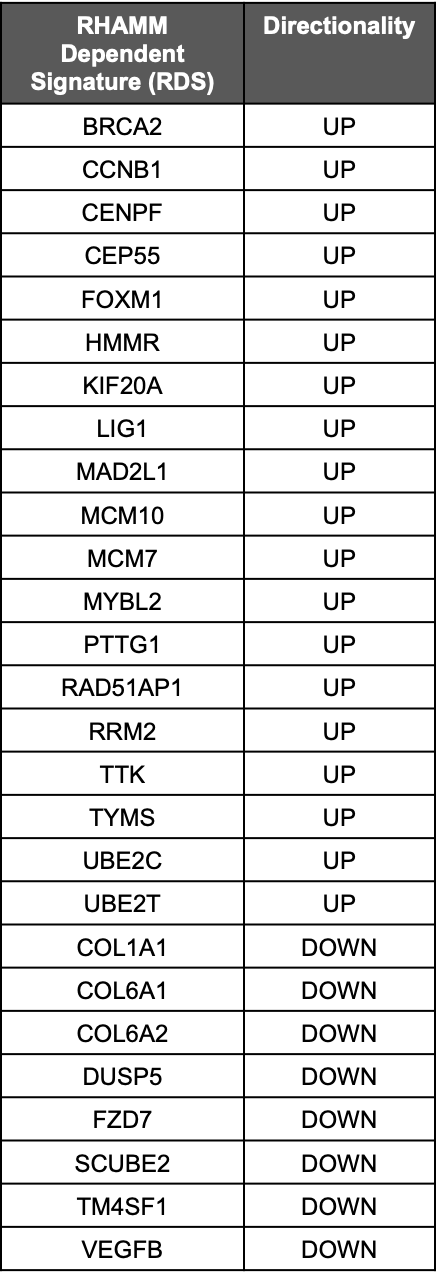

Supplemental Table 2. Enrichr pathway analysis of RDS

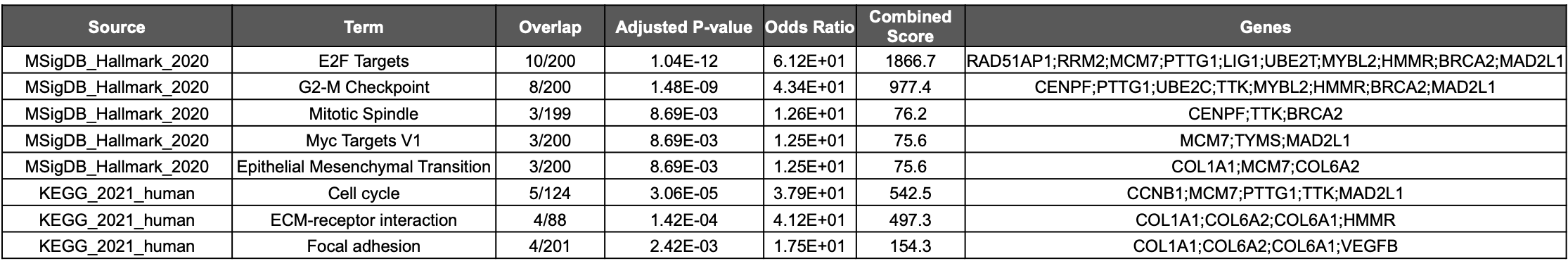

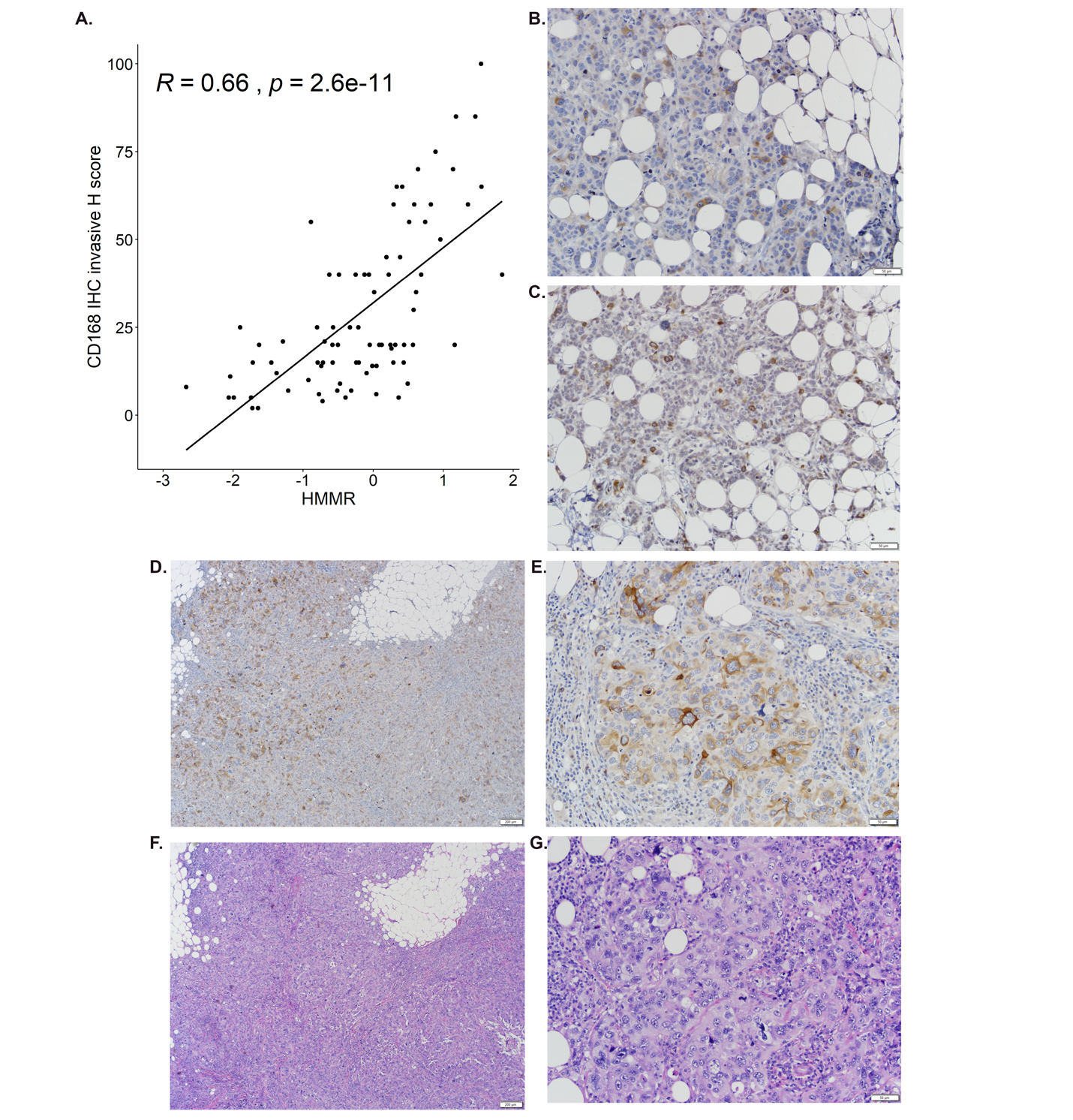
**Supplemental Figure 2. RHAMM expression is heterogeneous in human breast cancers.** (**A**) Correlation plot of *HMMR* transcript expression vs. RHAMM (CD168) protein H-score. (**B&C**) Representative images of RHAMM positive cells at invasive margins infiltrating into adjacent adipose tissue. (**D-G**) Representative image of high proportions of RHAMM+ cells at infiltrating margins of a high-grade tumor at low (**D**) and high (**E**) magnification, with (**F&G**) corresponding serial H&E stain.

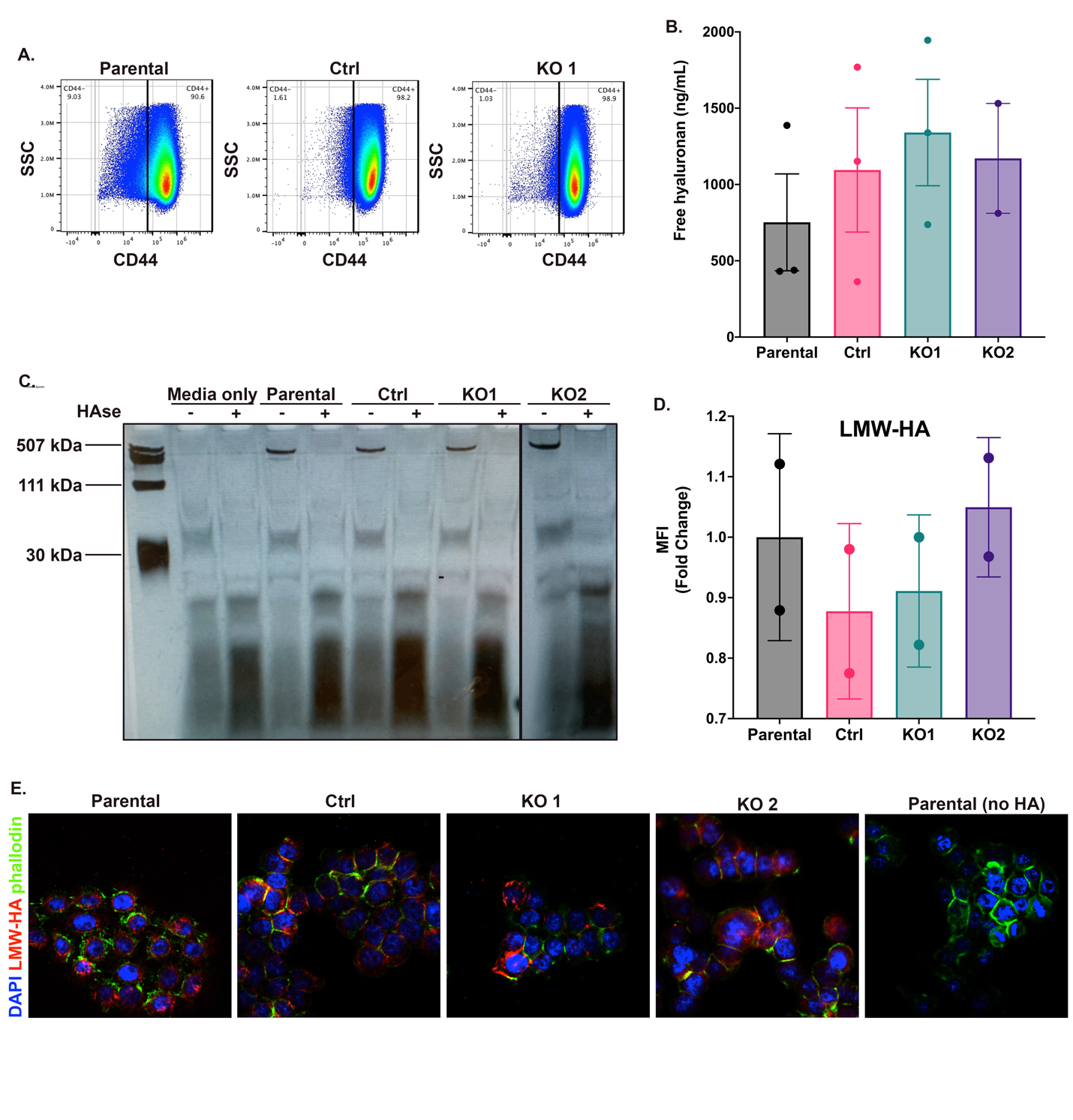
**Supplemental Figure 3. Characterization of RHAMM KO cell lines** (**A**) Representative image and analysis of cell surface CD44 expression via flow cytometry. (**B**) Extracellular HA concentration was determined in conditioned media from MCF10DCIS.com parental, control and RHAMM KO cell lines via an HA ELISA. (**C**) HA-fragmentation gel of conditioned media from MCF10DCIS.com parental, Ctrl, and RHAMM KO cell lines**,** hyaluronidase (HAse) treated conditions included as a control. (**D**) Mean fluorescent intensity (MFI) of labeled low molecular weight-HA (LMW-HA) treated MCF10DCIS.com parental, control and RHAMM KO cell lines. (**E**) Representative images of fluorescently labeled (LMW-HA) that has been taken up by MCF10DCIS.com parental, control and RHAMM KO cells.

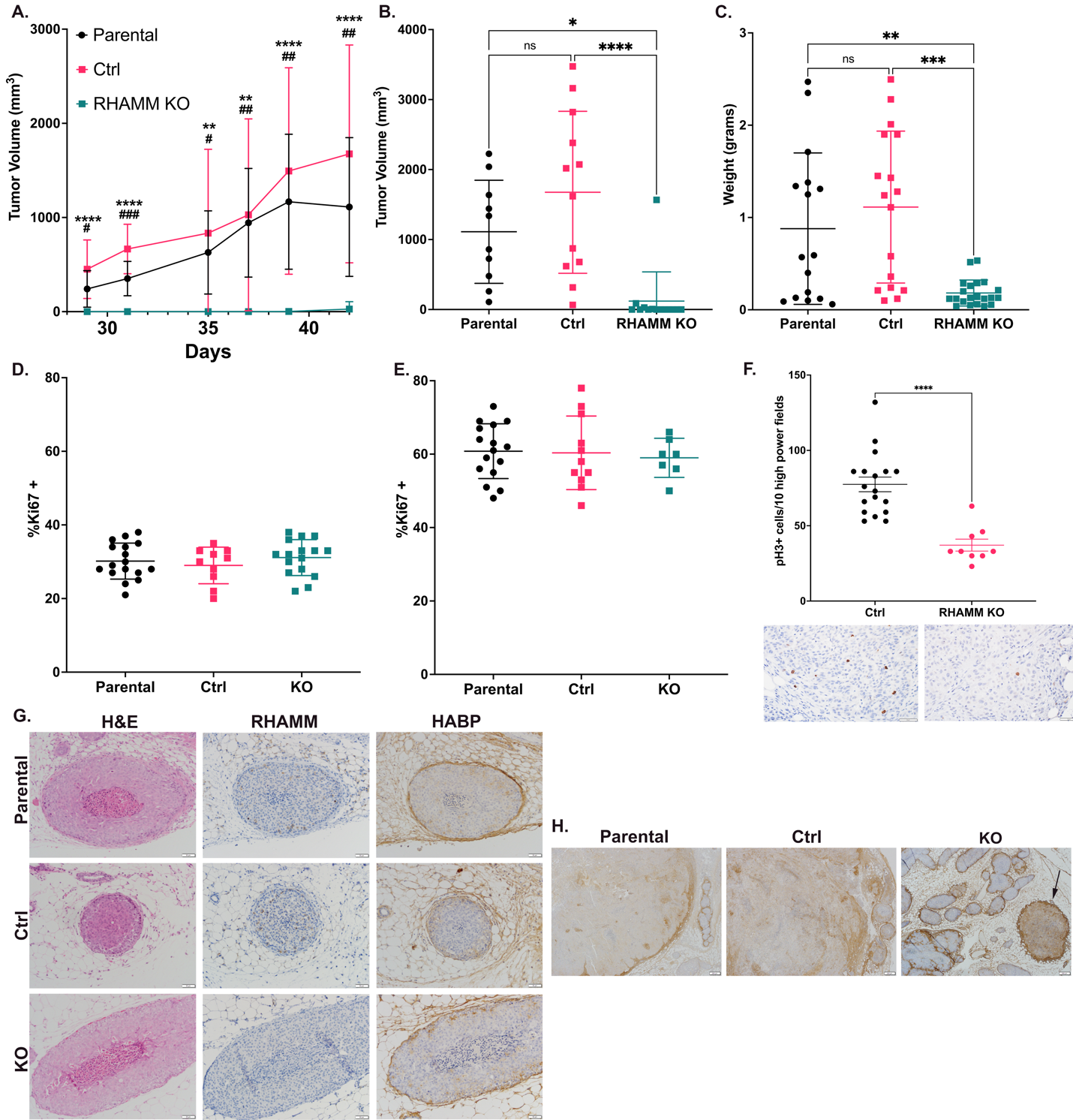

**Supplemental Figure 4. RHAMM drives invasion in vivo.** (**A**) Tumor volumes for mice injected with MCF10DCIS.com parental, control (Ctrl), and RHAMM KO cells, * T-test Parental vs KO, # T-tests Ctrl vs KO. (**B**) Final tumor volumes and **C.** mammary gland plus tumor weights for MCF10DCIS.com parental, control and RHAMM KO cells. (**D&E**) Ki67 IHC for *in situ* (**D**) or invasive (**E**) parental, Ctrl, or KO tumors. (**F**) Quantification of pH3 IHC for Ctrl and KO tumors, representative images below. (**G**) Representative images of *in situ* tumors stained for H&E and IHC for RHAMM and HABP. (**H**) Low power (40X) images of HABP staining shows increased intratumoral HA accumulation in invasive components (black arrow) of KO tumors compared to parental or control tumors. In situ components do not show the same accumulation of HA in the KO tumors.

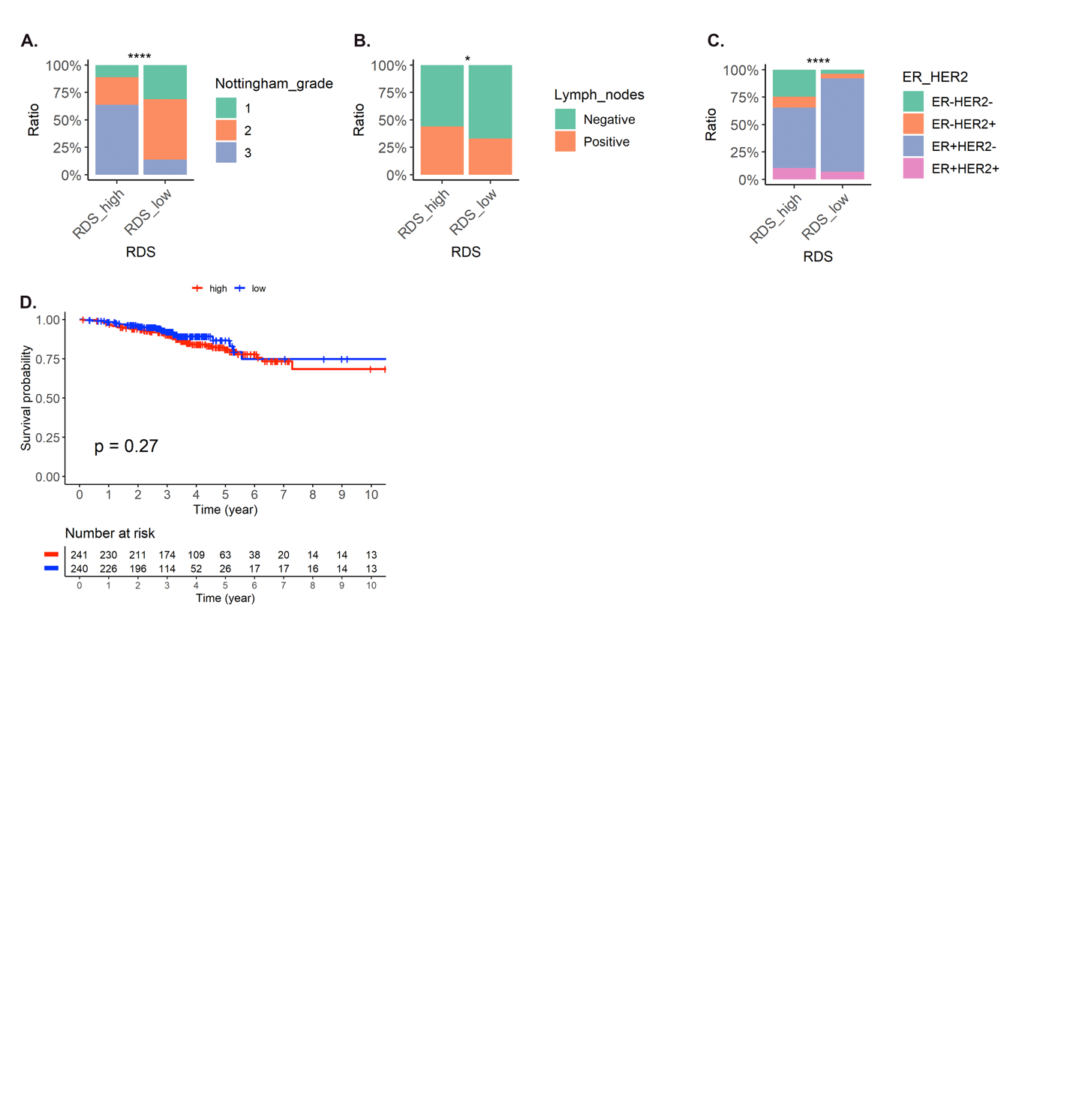

**Supplemental Figure 5**. **RDS correlates with poor outcomes in breast cancer** Distribution of clinicopathological features across RDS clusters of breast cancer patients from the BCCR, including Nottingham grade (**A**), lymph node (**B**), and clinical hormone receptor status (**C**). (**D**). Kaplan-Meier plot of BCCR cases by *RHAMM* high or low for overall survival at 10 years. * p value < 0.05, **** p value < 0.0001.

| **Antibody** | **Species** | **Dilution** | **Catalog NO.** | **Vendor** |
| --- | --- | --- | --- | --- |
| RHAMM, monoclonal | Rabbit | 1:1000 WB  1:100 IHC,IF | EPR4055 | Abcam Inc |
| β-tubulin, monoclonal | Rabbit | 1:5000 | 2146S | Cell Signaling Technology |
| Vimentin, monoclonal | Rabbit | 1:1000 | 5741S | Cell Signaling Technology |
| Anti-rabbit IgG, HRP-linked | Goat | 1:5000 | 7074S | Cell Signaling Technology |
| Ki67 | Rabbit | 1:400 | RM-9106-S1 | ThermoScientific |
| pH3 | Rabbit | 1:100 | 9701S | Cell Signaling Technology |
| HA-binding protein (HABP), Biotinylated | Bovine | 1:50 | 80502-722 | MilliporeSigma |
| CD44 | Rat | 1:200 | 17-0441 | eBioscience |

**Supplemental Table 3.** Antibody information
